# Opsin gene evolution in amphibious and terrestrial combtooth blennies (Blenniidae)

**DOI:** 10.1101/503516

**Authors:** Fabio Cortesi, Karen L. Cheney, Georgina M. Cooke, Terry J. Ord

## Abstract

Evolutionary adaptations to life on land include changes to the physiology, morphology and behaviour of an animal in response to physical differences between water and air. The visual systems of amphibious species show pronounced morphological adaptations; yet, whether molecular changes also occur remains largely unknown. Here, we investigated the molecular evolution of visual pigment genes (opsins) in amphibious and terrestrial fishes belonging to the Salariini division of blennies (Blenniidae). We hypothesized that when conquering land, blenny opsins adapt – in terms of sequence variation and/or gene expression – to match both higher light intensities as well as the broader light spectrum. Using retinal transcriptomes in six species ranging from fully aquatic to fully terrestrial, we found very little variation in opsin gene sequences or gene expression between species. All blennies expressed a single rod opsin gene as well as two cone opsin genes sensitive to longer-wavelengths of light: *RH2A-1* (green-sensitive) and *LWS* (red-sensitive). They also expressed one or two short-wavelength-sensitive cone opsin genes (*SWS2Aα, SWS2Aβ;* blue-sensitive) in a phylogenetically inert manner. However, based on amino acid predictions, both SWS2A proteins confer similar peak spectral sensitivities and differential expression is therefore unlikely to be ecologically significant. Red-sensitivity is likely beneficial for feeding on algae and detritus, the main food source of Salariini blennies, and could be co-adapted to perceive visual displays in terrestrial species, which often use red dorsal fins to signal during aggressive disputes and courtship. Our data suggests that on the molecular level, the visual systems that evolved in aquatic blennies have been retained in species that have transitioned onto land.

## Introduction

The evolution of tetrapods provides a textbook example for the transition from water-to-land and occurred in the Devonian around 380 million years ago (MacIver et al. 2017). Interestingly, terrestrial or semi-terrestrial (amphibious) lifestyles, have evolved independently in at least thirty-three osteichthyan families (Ord and Cooke 2016, Wright and Turko 2016). Selective forces that drive fishes to leave the water may include access to food, new or better resources for reproduction (Shimizu et al. 2006), an escape from aquatic predation (Ord et al. 2017) or reduced competition (reviewed in Sayer 2005, Wright and Turko 2016). However, plastic and/or evolutionary adaptations are often needed in response to the extreme environmental changes between water and air. Such adaptations to life on land include: aerial respiration (Mandic et al. 2009, Regan et al. 2011), mucus secretion to prevent dehydration (Smith 1930, Sturla et al. 2002), and shifts in ammonia household to avoid NH3 intoxication (Davenport and Sayer 1986, Chew and Ip 2014). Skeletal modifications are also common and may be triggered due to changes in locomotion (Kawano and Blob 2013, Brunt et al. 2016), an increase in body weight due to increase in apparent gravity (Turko et al. 2017), and by elevated oxygen content (Rossi et al. 2018). Moreover, behavioural modifications may occur, such as shade seeking to avoid desiccation during low tide (Colombini et al. 1995, Ord and Tonia Hsieh 2011).

The sensory systems of animals may also change as they move on to land. Since most fishes lack eyelids or similar structures, many amphibious fish species have modifications such as pigmented corneas and lenses, horizontal pupils and retractable eyes, to cope with increased light intensity on land (Sayer 2005). The refractive index of air versus water is also drastically different requiring modifications of the cornea, the lens or even the retina itself to maintain acuity when switching between environments (Sayer 2005). An extreme example of this can be found in the four-eyed fish, *Anableps anableps*, which has horizontally divided eyes: the top half has a flatter cornea compared to the bottom half to allow simultaneous vision above and below water, respectively (Swamynathan et al. 2003). However, while morphological adaptations have been studied in some detail, very little is known about possible molecular changes to the visual systems of amphibious and terrestrial fishes.

At the core of animal vision lie the opsin proteins that, together with a vitamin A-derived chromophore, form a light-sensitive photopigment located in the photoreceptors of the retina (Wald 1968, Hunt et al. 2014). Ancestral opsin gene duplications and subsequent changes to their amino acid compositions have led to five classes of vertebrate visual opsins that can be defined by their photoreceptor specificities and the different wavelengths of light they are maximally sensitive to (λmax) (Hunt et al. 2014). RH1, the rod opsin is expressed in rod cells and used for dim-light vision, while the four other opsin classes are expressed in cone photoreceptors and used for daylight colour vision. These are: two short-wavelength, ultraviolet (UV) and blue-sensitive opsins (SWS1 and SWS2); one medium-wavelength-sensitive green opsin (RH2); and a long-wavelength-sensitive red opsin (LWS) (Yokoyama 2002). In teleost fishes and amphibians with different cone morphologies, SWS opsins are further expressed in single cones, while RH2 and LWS opsins are found in double cones (two single cones that are fused together) (Hunt et al. 2014).

The evolution of visual opsins in teleost fishes is especially dynamic involving additional gene duplications, deletions, pseudogenizations and gene conversions, which has led to fishes having between one and 40 opsin genes in their genomes (Cortesi et al. 2015, Lin et al. 2017, Musilova et al. 2018 preprint). This diversity is primarily thought to be due to the different light environments fishes inhabit. For example, light at either end of the visible spectrum gets absorbed with increasing depth and consequently deeper living species gradually lose their *SWS1* and *LWS* genes (Musilova et al. 2018 preprint). Species living in murky red-light dominated habitats, on the other hand, often have red-shifted visual systems (e.g., Hofmann et al. 2009, Escobar-Camacho et al. 2016); however, this could also be due to sexual selection (see Sandkam et al. 2018 for a recent review on guppy *LWS* evolution). Mutations in the coding sequence, differential expression of opsin genes and switches between A1-based and A2-based chromophores also enable adaptations to more subtle differences in photic environments, such as between seasons (Shimmura et al., 2017), microhabitats (Fuller et al. 2010, Luehrmann et al. in review), feeding habits (Stieb et al. 2017), predation pressure (Sandkam et al., 2018), or in response to sexual selection (Seehausen et al. 2008).

In this study, we focused on the evolution of opsin genes in amphibious and terrestrial combtooth blennies (Blenniidae). Combtooth blennies are small (< 10 cm) scaleless fish that are commonly found in many shallow tropical and warm water marine habitats, including coral reefs, estuaries, mangroves, tide pools, and sometimes on land (Hundt et al. 2014a, Hundt and Simons 2018). They comprise one of the most diverse percomorph families consisting of 400 described species (58 genera; fishbase.org) that fall into 13 phylogenetic clades (Hundt et al. 2014a, Hundt and Simons 2018). Within the Salariini division of blennies, amphibious behaviour is common, and more than twenty species in at least three genera exhibit a highly terrestrial lifestyle (Ord and Cooke 2016). In these species, post-settlement larvae (∼30 days from hatching; Platt and Ord 2015) are believed to transition to a terrestrial lifestyle within the supralittoral splash zone and do not voluntarily return to the aquatic environment (Ord et al. 2017). Terrestrial blennies move freely about the rocks using a tail-twisting behaviour that allows them to efficiently shuffle along the rocks and jump distances of several body lengths (Hsieh 2010). Like most intertidal blennies, amphibious and terrestrial species feed primarily on detritus and algae (Hundt et al. 2014a, Hundt et al. 2014b, Hundt and Simons 2018) and possess a cryptic body colouration and patterning that reduces predation (Morgans and Ord 2013, Ord et al. 2017). However, unlike aquatic and most amphibious blennies, adults of terrestrial species in both sexes display a brightly coloured red-orange dorsal fin during aggressive disputes and courtship (Bhikajee and Green 2002, Shimizu et al. 2006, Ord and Tonia Hsieh 2011). These conspicuous fins contrast against the rocky backgrounds on which the blennies are active (Morgans and Ord 2013, Morgans et al. 2014), and are further accentuated in courting males by a darkening of the body to a largely uniform charcoal black (Ord and Tonia Hsieh 2011).

Morphological adaptation for aerial vision amongst the Salariini has been reported for the amphibious (possibly exclusively terrestrial) Kirk’s blenny, *Alticus kirkii*, which, by separating the cornea conjunctiva and cornea propria, has formed an additional eye chamber that is adjustable to accommodate changes in refracting indices of media (Zander 1974). Also, the retina of the mildly amphibious rippled rockskipper, *Istiblennius edentulus*, (see Ord and Cooke 2016) shows prominent swellings and folds, and a central depression into which the lens can be retracted. This allows light to be focused onto the back of the eye in both water and air (Zander 1974).

In terms of the molecular basis for vision, little is known about the evolution of opsin genes in Salariini blennies. However, a close relative to the Salariini, the rock-pool blenny, *Parablennius parvicornis*, was recently found to possess six cone opsins and one rod opsin in its genome (Musilova et al. 2018 preprint). Also, transcriptome sequencing in another aquatic blenny, the bluestriped fangblenny, *Plagiotremus rhinorhynchos*, revealed that this species expresses five orthologous cone opsins in its eyes (Musilova et al. 2018 preprint). Given that amphibious species show morphological adaptations for vision in air, and that blenny visual opsins are extremely diverse, we hypothesized that aerial vision and exposure to full sunlight caused adaptations in the Salariini opsin gene repertories. More specifically, due to the tissue damage UV-radiation inflicts, we expected to see a lack of *SWS1* expression in the eyes of these fishes. Instead, feeding on detritus and algae (Hundt et al. 2014b), which generally show strong reflection in the red due to their chlorophyll component (Stieb et al. 2017), as well as the use of red dorsal fins for aggressive and courtship displays (see Ord and Tonia Hsieh 2011, Morgans and Ord 2013), might have caused amphibious and terrestrial blennies to have a more long-wavelength, *LWS*, dominated visual system.

## Material and Methods

### Study species

Adult blennies belonging to the Salariini clade (Hundt et al. 2014a) were collected on snorkel or on foot with hand nets from several sites around the Indo-Pacific. Given that these blennies actively seek shade to avoid desiccation (Ord and Tonia Hsieh 2011), sampling either occurred in the early hours of the morning or late in the afternoon. Immediately post capture, their eyes were enucleated and stored on RNAlater (Life Technologies) for subsequent molecular analysis. From French Polynesia (FP), we collected Lined Rockskipper, *Istiblennius lineatus* (Mo’orea: 17°29′52.05ʺS, 149°45′17.24ʺW; November 8-10, 2013; N = 3), Blackmargin Rockskipper, *Praealticus caesius* (Mo’orea: 17°29′52.05ʺS, 149°45′17.24ʺW; November 8-10, 2013; N = 3); and Marquesan Rockskipper*, Alticus simplicirrus* (Tahiti: 17°30′57.67ʺS, 149°24′25.14ʺW; November 14-15, 2013 N = 3). From the Seychelles (S), we collected *I. lineatus* (4°48′9ʺS, 53°30′60ʺE and 4°33′56ʺS, 55°27′10ʺE; April 19 and 26, 2014; N = 3); Reef Margin Blenny, *Entomacrodus striatus* (4°48′9ʺS, 53°30′60ʺE; April 19 and 22, 2014; N = 3); and the Seychelles Rockskipper, *Alticus anjouanae* (4°33′56ʺS, 55°27′10ʺE; April 20, 2014; N = 3). A Jewelled Blenny, *Salarias fasciatus* (N = 1), was collected in May 2017 from Heron Island (23°44′S, 151°91′E), Great Barrier Reef, Australia.

All experimental procedures were approved by Animal Ethics Committees from The University of New South Wales (11/36B and 13/21) and The University of Queensland (QBI/304/16). Fish were collected under permits issued by Protocole D’Accueil (10/10/2013) French Polynesia, Seychelles Bureau of Standards (#A0157), the Great Barrier Reef Marine Park Authority (G17/38160.1) and Queensland Fisheries (#180731).

### Transcriptome sequencing, quality filtering and de-novo assembly

Retinae were dissected out of the eyecup and total RNA was extracted using an RNAeasy Mini Kit (Qiagen) including DNAse treatment following the manufacturer’s instructions. RNA integrity was checked using an Eukaryotic Total RNA Nano chip on a Bioanalyzer 2100 (Agilent Technologies). All retinal transcriptomes, except for *S. fasciatus*, were sequenced in-house at the Queensland Brain Institute’s sequencing facility, Brisbane, Australia. Sequencing libraries were prepared from 100-1,000 ng of total RNA using the TruSeq total stranded mRNA Library Prep Kit protocol (Illumina, San Diego), and library concentrations were measured using a Qubit dsDNA BR Assay Kit (Thermo Fisher). Individual libraries were barcoded and up to 12 libraries/lane were pooled at equimolar ratios. Libraries were sequenced at PE125 on a HiSeq 2000 using Illumina’s SBS chemistry version 4. The *S. fasciatus* library preparation (strand-specific, 250∼300 bp insert) and sequencing (RNAseq HiSeq PE150) was outsourced to Novogene (https://en.novogene.com/).

Filtering and *de novo* assembly of retinal transcriptomes followed the protocol described in (de Busserolles et al. 2017). In short, raw-read transcriptomes were uploaded to the Genomics Virtual Laboratory (GVL 4.0.0) (Afgan et al. 2015) on the Galaxy Australia server (https://galaxy-qld.genome.edu.au/galaxy/) and quality filtered using Trimmomatic (Galaxy v.0.32.2) (Bolger et al. 2014) before being *de novo* assembled using Trinity (Galaxy v.0.0.2) (Haas et al. 2013). Two transcriptomes per species were assembled, with the exception of *S. fasciatus* where only one individual was sequenced. Raw-read libraries and assemblies are available on the Sequence Read Archive (https://www.ncbi.nlm.nih.gov/sra) and the Transcriptome Shotgun Assembly Database (https://www.ncbi.nlm.nih.gov/genbank/tsa/), respectively (Table S1).

### Opsin gene mining, phylogenetic reconstructions and opsin gene expression

Opsin gene mining and expression analyses followed the protocol described in de Busserolles et al. 2017. In short, putative Salariini opsin genes were searched for by mapping the assembled transcripts to the opsin coding sequences of the dusky dottyback, *Pseudochromis fuscus* (Cortesi et al. 2016) in Geneious v.11.0.2 (www.geneious.com). *Ps. fuscus* was chosen because it belongs to the closely related Pseudochromidae (Alfaro et al. 2018) and possesses an opsin gene repertoire containing representatives from all ancestral vertebrate opsin gene classes (Cortesi et al. 2016). Mapped contigs were extracted and compared to publicly available opsin gene sequences using BLASTN (https://blast.ncbi.nlm.nih.gov/Blast.cgi). Moreover, because *de novo* assembly based on short-read libraries is prone to create misassemblies (chimeric sequences), and/or to overlook closely related or lowly expressed gene copies, we used a second approach to account for all expressed Salariini visual opsins. This approach entailed the mapping of unassembled reads to the reference *Ps. fuscus* genes, followed by a manual extraction of gene copies using paired-end information to move from single polynucleotide polymorphism (SNP) to SNP along the gene. Extracted reads were *de novo* assembled and if necessary, their consensus was used as a species-specific template against which unassembled reads were repeatedly re-mapped until the whole coding region could be extracted.

Salariini visual opsins were confirmed and assigned to specific opsin gene classes based on their phylogenetic relationships to a reference dataset obtained from GenBank (https://www.ncbi.nlm.nih.gov/genbank/) and Ensembl (www.ensembl.org/) (Fig. 1, Fig. S1). Gene coding regions were aligned using the L-INS-I settings as part of the Geneious MAFFT plugin v.1.3.7 (Katoh and Standley 2013), and jModeltest v.2.1.10 (Darriba et al. 2012) was subsequently used to select the most appropriate model of sequence evolution based on the Akaike information criterion. MrBayes v.3.2.6 (Ronquist et al. 2012) run on the CIPRES platform (Miller et al. 2010) was then used to infer the phylogenetic relationship between opsin genes using the following parameter settings: GTR+I+G model; two independent MCMC searches with four chains each; 10 million generations per run; 1000 generations sample frequency; and, 25% burn-in. Phylogenies were also reconstructed using GTR+G models, to account for the possibility of variable sites, however, no substantial differences in tree structure or node support could be found. Raw trees from either approach and corresponding data alignments have been deposited in Dryad (https://datadryad.org/), and GenBank accession numbers for the relevant genes are given either in Fig. S1, or in Table S1.

**Fig 1.**
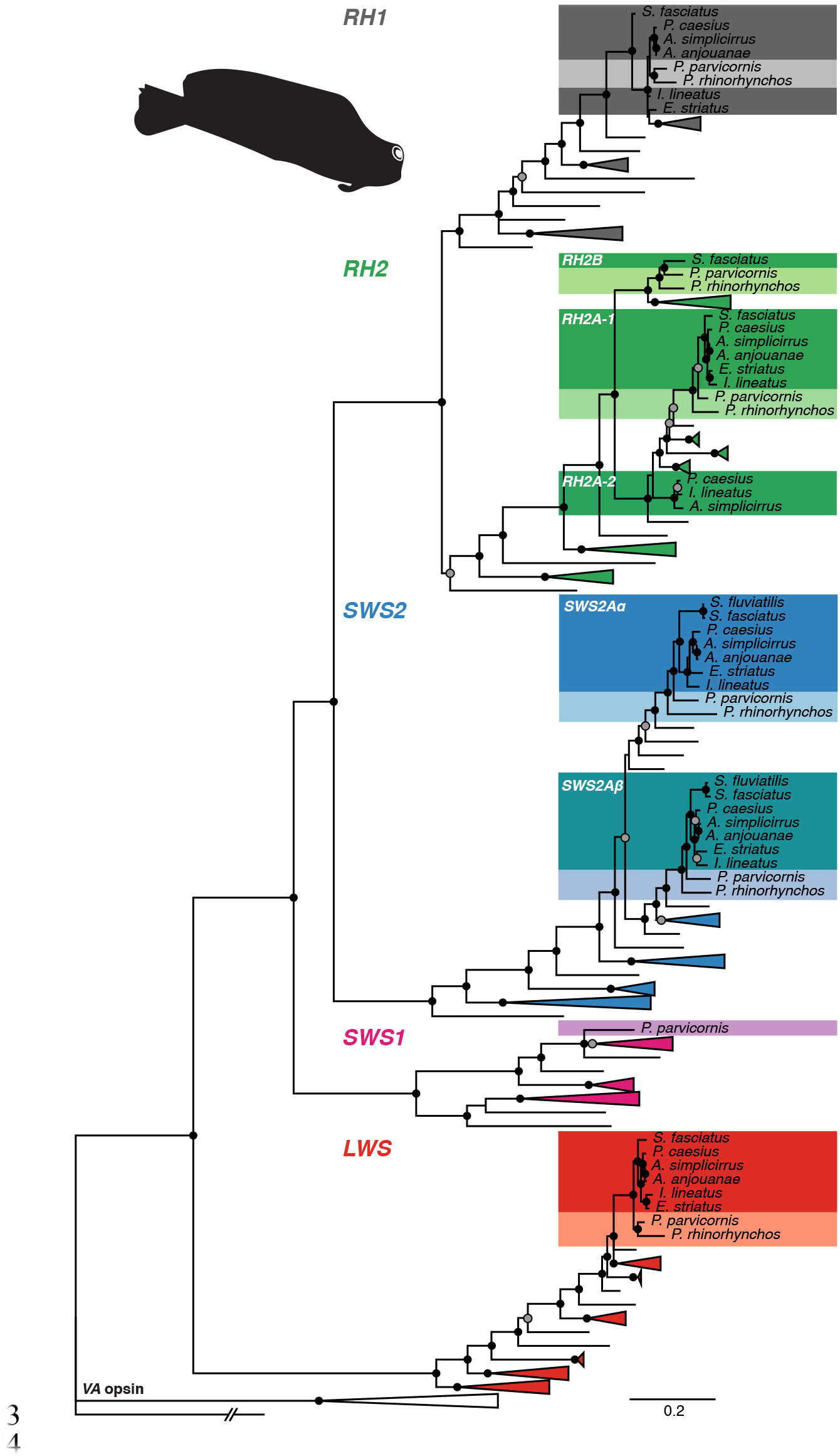
Bayesian consensus phylogeny for vertebrate opsin genes. The Salariini retinal transcriptomes contained seven opsin genes. One rod opsin (*RH1*) and six cone opsin genes belonging to three different cone opsin classes: short-wavelength-sensitive 2 (*SWS2Aα, SWS2Aβ*), rhodopsin-like 2 (*RH2A-1, RH2A-2, RH2B*), and long-wavelength-sensitive (*LWS*). Note that *RH2A-2* was found at very low expression levels in three out of the six Salariini species, while *RH2B* was lowly expressed in *Salarias fasciatus* alone (also see Fig. 2, Table 2). Dark shading indicates genes from various Salariini species, light shading genes from sister species (Blenniidae). Black and grey spheres indicate Bayesian posterior probabilities > 0.9 and 0.7, respectively.

Quantitative opsin gene expression was measured by mapping the unassembled reads against the extracted opsin coding regions for each species as per de Busserolles et al. 2017. We then compared the rod opsin expression to the combined cone opsin expression, the proportional expression of each cone opsin gene to the combined cone opsin expression and finally, the proportional expressions of single (*SWS2Aα, SWS2Aβ*) and double cone (*RH2A-1, LWS*) genes amongst themselves (Table 2). Single and double cone opsin gene expression for each species was then plotted onto the Salariini phylogeny taken from Ord and Cooke, 2016 (Fig. 2). Moreover, we also quantified the expression of *cytochrome P450 family 27 subfamily c member 1 (cyp27c1)* by comparing its gene expression to the total opsin expression of fishes. The zebrafish *cyp27c1* ortholog converts vitamin A1-based chromophore to the longer shifted A2-chromphore (Enright et al. 2015) and A2 has previously been reported in the peacock blenny, *Salaria pavo*, based on microspectrophotometry results (White et al. 2004).

**Fig 2.**
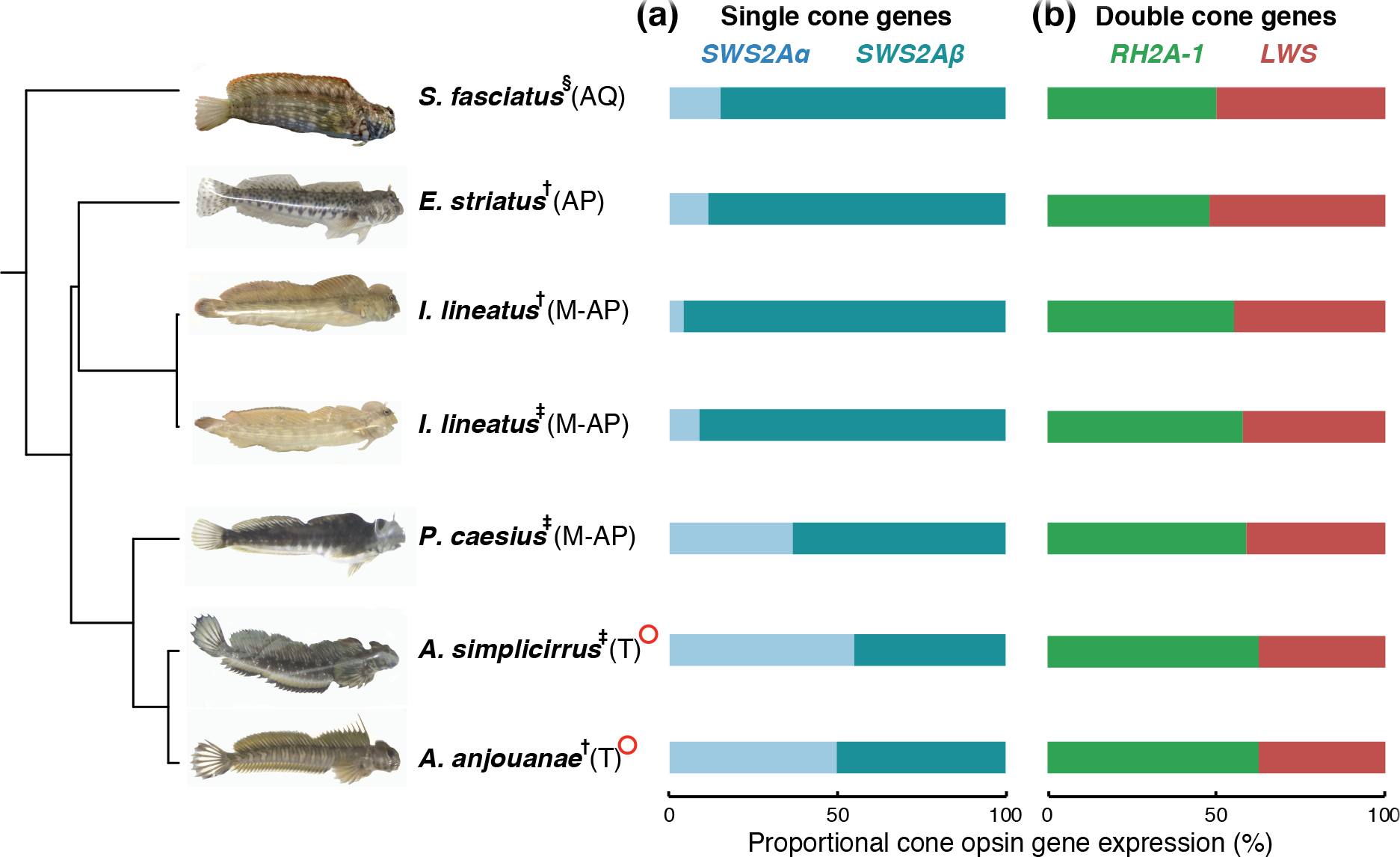
Salariini species phylogeny and associated opsin gene expression. a) Salariini species (n = 3 per species, except for *S. fasciatus* where n = 1) expressed two paralogs of the short-wavelength-sensitive 2A opsin gene (*SWS2Aα, SWS2Aβ*), and b) one rhodopsin-like 2 (*RH2A-1*) and a long-wavelength-sensitive (*LWS*) opsin gene. The mean proportional gene expression was similar between species independent of habitat or sampling location, except for the *SWS2A* paralogs in the clade containing *Pr. caesius* and the two *Alticus* spp. Displayed are the mean expression values separated by cone specificity (Hunt et al., 2014). For details on individual expression values and gene expression in relation to total cone and total opsin expression see Table 2. A red circle indicates terrestrial *Alticus* spp., which use red dorsal fins for aggressive and courtship displays (Bhikajee and Green 2002, Shimizu et al. 2006, Ord and Tonia Hsieh 2011). AQ = aquatic, AP = amphibious, M-AP = mildly-amphibious, T = Terrestrial, definitions and phylogeny as per (Ord & Cooke, 2016). † Seychelles, ‡ French Polynesia, § Heron Island (Australia).

### Opsin gene sequence analysis and spectral sensitivity predictions

Opsin gene coding sequences were aligned using the L-INS-I settings in MAFFT (Geneious plugin v.1.3.7) (Katoh and Standley 2013) and gene specific opsin trees were reconstructed using RAxML v.8.2.11 (Geneious plugin v.3.0) (Stamatakis 2014), a GTR+G model and 1000 bootstraps to generate the support values for majority-rule consensus trees (Fig. S2). The alignments and corresponding trees were then used to test for site-specific positive selection using codeml in PAML (Yang 2007) as described in detail by (Hofmann et al. 2012). Briefly, codeml was run on the graphical user interface pamlX v.1.3.1. (Xu and Yang 2013) using likelihood ratio tests (LRT) to compare M1a vs. M2 and M8 vs. M8a. Bayesian Empirical Bayes (BEB) criteria (Yang et al. 2005) were subsequently applied to identify specific amino acid sites under positive selection in case of significant LRTs (Table S2). Next, we imported the alignments into MEGA7 (Kumar et al. 2016), which was used to calculate synonymous (ds) and nonsynonymous substitution rates (dn) using the Nei-Gojobori method (Jukes-Cantor distances)(Table 1).

**Table 1.** Summary of Salariini opsin gene variation

|  | *SWS2Aα* | *SWS2Aβ* | *RH2A-1* | *LWS* | *RH1* |
| --- | --- | --- | --- | --- | --- |
| Total number of nucleotides | 1053 | 1056 | 1059 | 1074 | 1065 |
| Total number of amino acids | 350 | 351 | 353 | 357 | 354 |
| Variable nucleotide sites | 140 | 105 | 64 | 71 | 85 |
| Indels | 0 | 0 | 0 | 0 | 0 |
| <u>Between Salariini species (amino acids)</u> |  |  |  |  |  |
| Variable transmembrane sites | 13(6) | 10(2) | 3(2) | 10(3) | 11(3) |
| Variable retinal binding pocket sites | 1(0) | 1(0) | 0 | 1(1) | 2(0) |
| N° of substitutions at known opsin tuning sites | 0 | 4(0*) | 0 | 4(1*) | 1(1*) |
| Nonsynonymous substitutions (*dn*) | 0.014 | 0.009 | 0.003 | 0.006 | 0.008 |
| Synonymous substitution (*ds*) | 0.205 | 0.147 | 0.093 | 0.107 | 0.110 |
| *dn/ds* | 0.068 | 0.061 | 0.032 | 0.056 | 0.073 |
| <u>vs. *Parablennius parvicornis* (amino acids)</u> |  |  |  |  |  |
| Variable transmembrane sites | 22(10) | 18(4) | 10(2) | 12(3) | 14(4) |
| Variable retinal binding pocket sites | 2(1) | 1(0) | 1(0) | 1(1) | 3(1) |
| Number of substitutions at known tuning sites | 1(0*) | 5(0*) | 0 | 5(1*) | 2(2*) |

( ) substitution with changes in polarity; \* substitutions at known tuning sites of gene in question.

Salariini opsin amino acid sequences were then aligned to bovine rhodopsin (Palczewski et al. 2000) to assess their variability within the transmembrane and the retinal binding pocket sites as well as at known opsin spectral tuning sites (Fasick and Robinson 1998, Hunt et al. 2001, Yokoyama 2008, Dungan et al. 2015). Since the dn/ds ratio indicated low variability between Salariini opsins, we extended this analysis to also include the opsins of the rock-pool blenny, *Parablennius parvicornis* (Musilova et al. 2018 preprint), as a more distantly related obligate aquatic species from the sister clade of the Parablenniini (Hundt et al. 2014a). Amino acid substitutions were assessed for each opsin gene amongst Salariini species, and between the Salariini and *Pa. parvicornis* taking specific note of sites that differ in polarity between species (as per Hofmann et al. 2012) (Table 1 and Table S3). Finally, we inferred the spectral sensitivities of the Salariini opsins based on changes at known spectral tuning sites compared to the well documented opsin sequences of *Ps. fuscus* (Cortesi et al. 2016) (Table S3). Individual sites are referred to according to their location relative to bovine rhodopsin.

## Results

### Opsin gene phylogeny and gene expression

Retinal transcriptomes from our studied species contained six cone and one rod opsin (*RH1*) gene. Phylogenetic reconstruction revealed two blue-sensitive *SWS2A* copies (*SWS2Aα, SWS2Aβ*), three green-sensitive *RH2* copies (*RH2A-1, RH2A-2* and *RH2B*), and one red-sensitive *LWS* gene (Fig. 1, Fig. S1, Table S1).

Rod opsin expression was generally much higher than cone opsin expression (> 73% of total opsin expression), except in *E. striatus* and *I. lineatus* (S) where rod and cone opsin genes were more equally expressed (~ 50% of total opsin expression each). In terms of cone opsin expression, single cone genes (*SWS2Aα + SWS2Aβ*) accounted for ~ 10 – 15% of total cone opsin expression, except for one *Pr. caesius* individual which had 24.4 % single cone gene expression (Table 2). Comparing single and double cone expression separately, the single cone specific *SWS2A* paralogs were expressed at similar ratios (~ 40 – 60 % each) within the subclade containing the two terrestrial *Alticus* spp. as well as the immediate sister group to this genus, the mildly-amphibious *Pr. caesius*. In the remaining mildly-amphibious, amphibious and aquatic species, *SWS2Aβ* was primarily expressed (> 83%) (Fig. 2, Table 2). Neither the UV-sensitive *SWS1* nor the violet-sensitive *SWS2B* single cone genes were found to be expressed in the retinae of adult Salariini. Considering double cone gene expression, all blennies were found to express high amounts of *RH2A-1* and *LWS* (~ 40 – 60% each)(Fig. 2, Table 2). *RH2A-2*, on the other hand, was found to be lowly expressed (< 0.1% of total cone opsin expression) only allowing full coding sequence reconstruction for *A. simplicirrus* and partial sequence reconstructions for *Pr. caesius* and *I. lineatus* (Fig. 1). *S. fasciatus* was the only Salariini species with low levels of *RH2B* expression (< 0.1% of total cone opsin expression). Therefore, gene expression for *RH2A-2* and *RH2B* are unlikely to be relevant for adult Salariini vision and have been excluded from all further analyses (data not shown). *cyp27c1* was found to be lowly expressed in all blennies at < 0.5% compared to total opsin expression (Table 2).

**Table 2.**
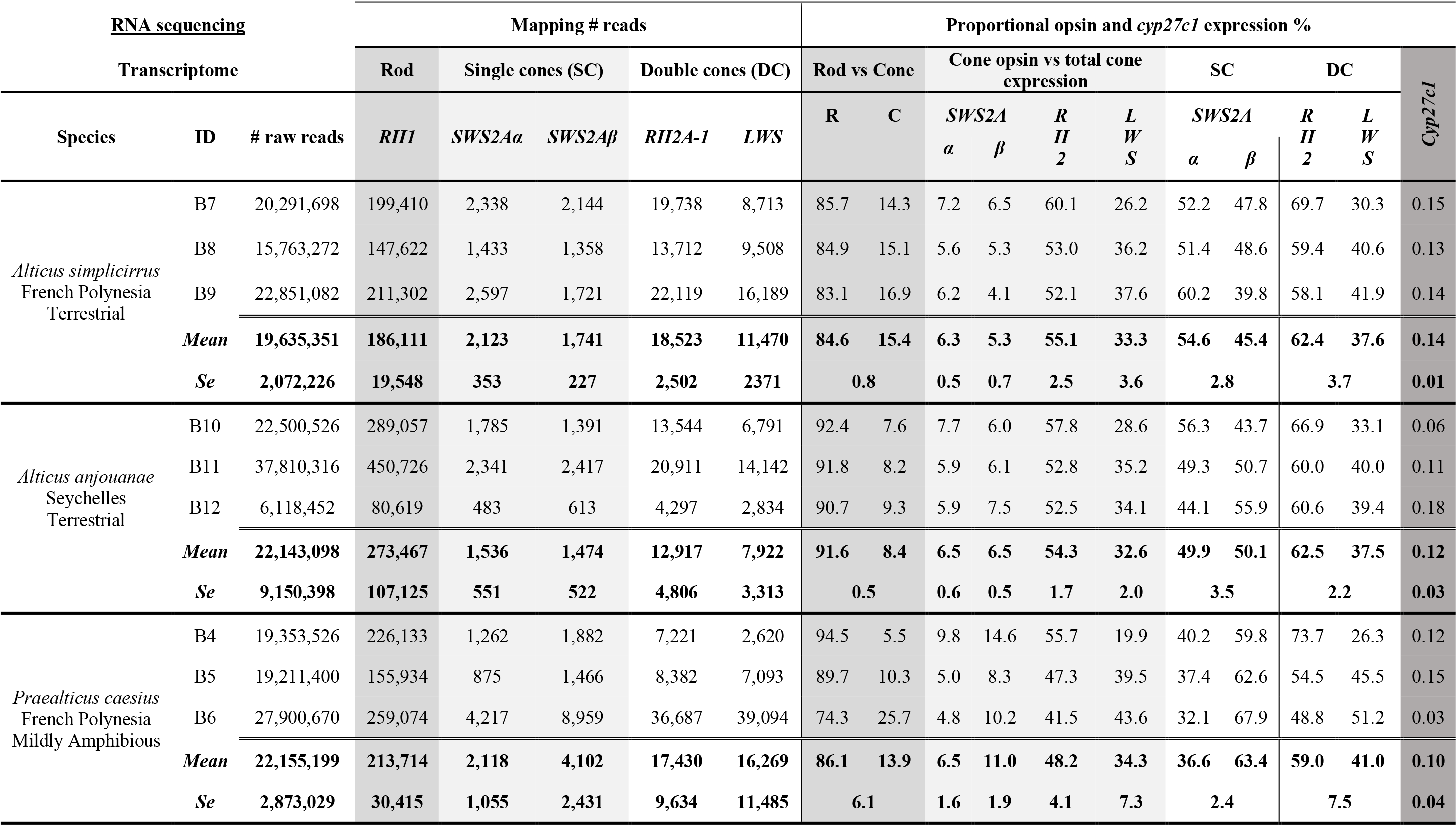
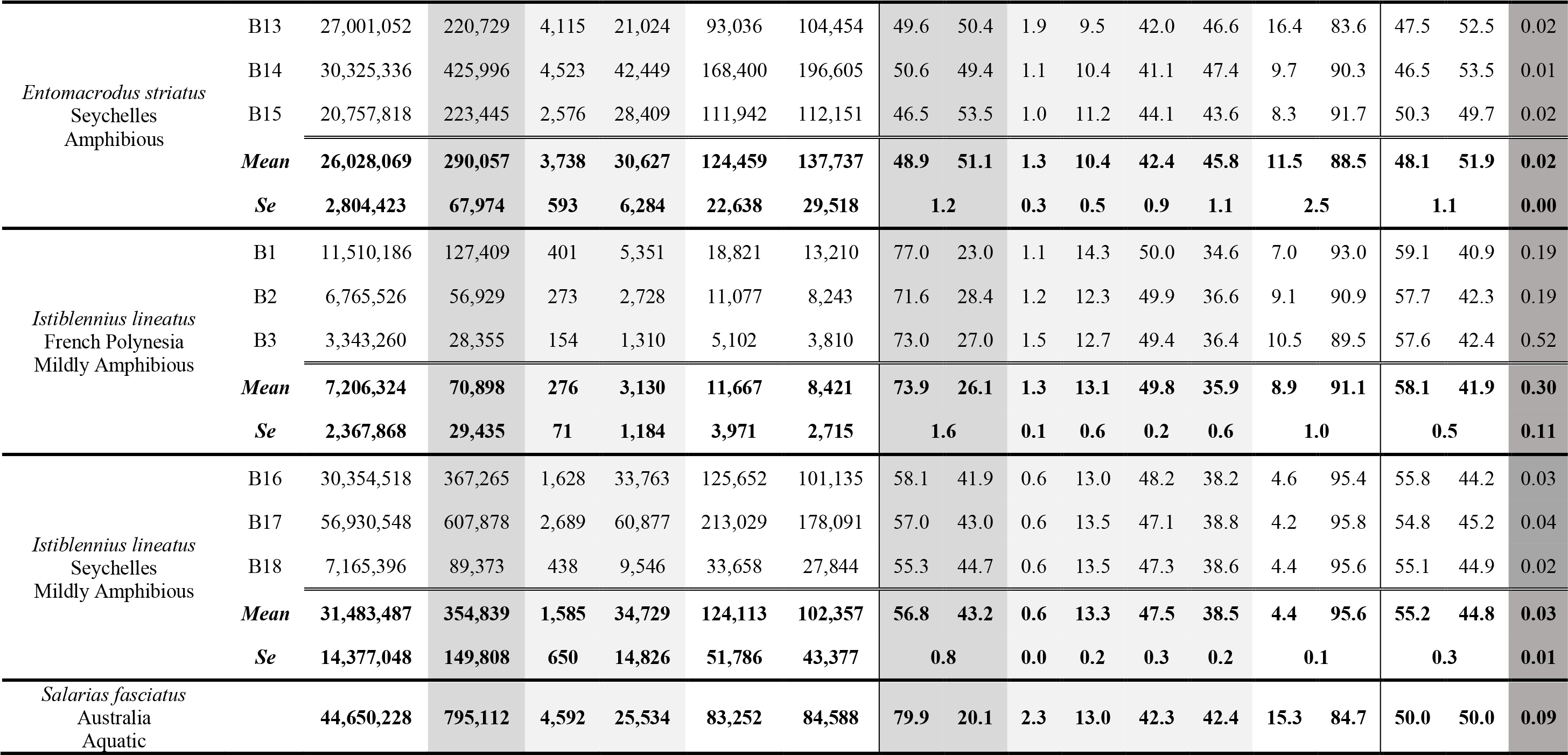
Summary of transcriptomes, opsin mapping, and proportional opsin gene expression. # raw reads refers to the total number of paired-end fragments. *RH1* = rod opsin, *SWS2* = short-wavelength sensitive 2, *RH2* = rhodopsin like 2, *LWS* = long-wavelength sensitive

### Opsin gene diversity, selection analysis and predicted spectral sensitivities

The Salariini opsin genes were highly similar both when compared between species within the clade, and when compared to the more distant *Pa. parvicornis* sister species (Table 1). Overall, *SWS2Aα* (13/22 amino acid substitutions within Salariini/against *Pa. parvicornis*) had the highest and *RH2A-1* (3/11) the lowest number of amino acid substitutions within transmembrane- and retinal binding pocket sites. Only a few of these amino acid substitutions were found to occur in potential spectral tuning sites, namely those sites that have previously been identified to change the spectral sensitivity of opsin pigments and/or substitutions that incorporated changes in polarity (Tables 1 and S3). Reflective of the low variability between the Salariini genes, using PAML revealed that none of the Salariini genes are under positive selection (Table S2).

Most Salariini opsin genes showed no changes in key tuning sites when compared to their *Ps. fuscus* orthologs (Cortesi et al. 2016; Table S3). Consequently, assuming a purely A1 chromophore based visual system, the peak spectral sensitivity (λmax) for the Salariini SWS2Aα was estimated to be at 448 nm. For SWS2Aβ we found one substitution at site T269A. The reverse substitution i.e. A269T, causes a 6 nm red-shift in λmax (Yokoyama 2008) and we therefore assumed a similar shift but in the opposite direction in our case. Thus, the Salariini SWS2AP was estimated to have a 451 nm λmax. The Salariini RH2A-1was found to be identical in key spectral tuning sites to the *Ps. fuscus* RH2Aα and its estimated λmax therefore to be 524 nm. The Salariini LWS (561 nm λmax) was also identical in tuning sites to its *Ps. fuscus* ortholog, expect for *P. ceaseus* (554 nm λmax) where the substitution S164A is likely to have blue shifted the spectral sensitivity by 7 nm (Yokoyama 2008). Finally, the Salariini RH1 (500 nm λmax) showed one substitution at N83D likely to cause a 2 nm red-shift (Yokoyama 2008) and in the case of the *S. fasciatus* RH1 (501 - 503 nm λmax), showed a second substitution at A124G further red-shifting this visual pigment by 1 - 3 nm (Hunt et al. 2001).

## Discussion

In this study, we investigated the molecular evolution of vision in fishes from the Salariini division of blennies, which have transitioned from water to land. First, we sequenced the retinal transcriptomes of six species classified as fully aquatic (*S. fasciatus*), mildly amphibious (*I. lineatus* and *Pr. caesius*), amphibious (*E. striatus*), and fully terrestrial (*A. simplicirrus* and *A. anjouanae*) (Ord and Cooke 2016; Fig. 1). We found that within their eyes, blennies express one rod opsin (*RH1*) and between three and four cone opsins (Fig. 2) independent of habitat or geographic region (i.e. South Pacific *vs.* Indian Ocean). *RH1* expression in fishes shows a strong diurnal pattern, starting with high expression in the morning and gradually decreasing as the day goes on (e.g., Korenbrot and Fernald 1989, Stieb et al. 2016). Hence, differences in sampling time i.e. early morning versus late afternoon, most likely explains the discrepancy of rod to cone opsin expression found in this study (Table 2).

Regarding the cone opsins, all Salariini species expressed the double cone genes sensitive to the red (*LWS*) and the green (*RH2A-1*) part of the light spectrum. We also found a second *RH2A* paralog in the genera *Alticus, Praealticus* and *Istiblennius* (Fig. 1). The *RH2A-2* gene was not expressed in the closely related *S. fasciatus* nor is it present in the genome of the more distantly related *Pa. parvicornis* (Musilova et al. 2018 preprint). One could therefore assume that this *RH2A* duplication is specific to the clade containing the amphibious and terrestrial Salariini species (Ord and Cooke 2016). However, based on our phylogenetic reconstruction, *RH2A-2* is basal to a greater *RH2A* clade which includes gene orthologs from dottybacks and cichlids (Fig. S1). Both cichlids (Escobar-Camacho et al. 2016) and dottybacks (Cortesi et al. 2016) have two *RH2A* duplicates which have undergone widespread gene conversion and consequently cluster closely together within species/families. The blenny *RH2A* copies, on the other hand, appear unaffected by gene conversion. Rather than being Salariini specific then, it is likely that *RH2A-2* is the blenny ortholog of an ancestral *RH2A* duplication and that *Pa. parvicornis* has lost the copy secondarily. As more and more fish genomes become available (e.g. Musilova et al. 2018 preprint), the patterns surrounding *RH2* evolution are likely to become clear in the near future.

Finally, there was a strong phylogenetic signal in the expression of the blue-sensitive *SWS2A* duplicates: only the clade containing the two *Alticus* species and *Pr. caesius* expressed both of the copies at a similar ratio, while the remaining Salariini species mainly expressed the slightly longer tuned *SWS2Aβ* copy (Fig. 2). Whether this difference in expression is ecologically significant or whether it is the result of a phylogenetic inertia remains to be investigated. Given that the predicted λmax for the two SWS2A paralogs is only 3 nm apart, we currently favour the latter scenario. As such, based on cone opsin expression and chromophore A_1_-derived spectral sensitivity predictions, Salariini blennies appear to have a well-developed, potentially trichromatic, colour vision sense which reaches across the visible light spectrum ranging from 448 - 561 nm in λmax.

Our study species live in shallow reef environments, are amphibious or have left the aquatic realm altogether. The light environment these species experience differs depending on their habitat with both an increase in light intensity and in the proportion of UV and red wavelengths when moving into shallower water and finally onto land (see e.g., Marshall et al. 2003). Opsin gene expression in fishes has been shown to be influenced by changes in light environment with water depth (e.g., Stieb et al. 2016), and degree of suspended organic material (e.g., Fuller et al. 2005). However, our data shows that there are no large differences in opsin gene expression between Salariini species from different habitats (Fig. 2; Table 2). All of the species appear to have red-shifted visual systems hinting towards the importance of detecting longer-wavelengths of light for survival. Mounting evidence suggests that long-wavelength reception in coral reef fishes is especially beneficial when feeding on algae or similar chlorophyll containing organic matter, which strongly reflect the red part of the light spectrum (for a recent review on the topic see Marshall et al. 2018). It is also possible that the red-orange dorsal fin that appears unique to the terrestrial species (TJ Ord, unpublished data) has evolved to exploit the pre-existing (ancestral) visual state of seeing red as a means of maximizing the efficiency of territorial and courtship signalling on land (Bhikajee and Green 2002, Shimizu et al. 2006, Ord and Tonia Hsieh 2011). The red-orange coloration of the dorsal fin of terrestrial blennies has been shown to be highly chromatically contrasting against the typical environmental background found on land (Morgans and Ord 2013). Similarly, sexually selected red-orange colour signals evolving in response to an innate red sensory bias in conspecific receivers has also been suggested for a number of other fish species (e.g., see Rodd et al. 2002, Smith et al. 2004, Seehausen et al. 2008).

It is notable that none of the species was found to express the UV-sensitive *SWS1* gene or the violet-sensitive *SWS2B* gene. While it is possible that *SWS2B* was lost in the ancestor of all blennies (Cortesi et al. 2015), *SWS1* is present, at least on the genomic level, in *Pa. parvicornis* (Musilova et al. 2018 preprint). Salariini blennies could have lost *SWS1* independently or alternatively, *SWS1* might simply not be expressed in adult blennies. Instead, *SWS1* and also the *RH2A-2* and *RH2B* paralogs, might be used at different developmental stages, a common feature of opsin gene expression in fishes (e.g., Spady et al. 2006, Carleton et al. 2008, Cortesi et al. 2016, Savelli et al. 2018). Supporting the potential use of *SWS1* at earlier life stages, Siebeck and Marshall (2007) found that adult Salariini blennies have UV blocking eyes while larval stages have UV to violet transmitting eyes. In an interesting parallel, mudskippers (Gobidae) (You et al. 2014) and the Asian swamp eel (*Monopterus albus*) (Musilova et al. 2018 preprint), both of which are amphibious, have lost their *SWS1* gene with the swamp eel further missing its *SWS2B* copy (Cortesi et al. 2015). It is possible that the tissue damage UV-radiation induces has led to the convergent loss and/or the inexpression of the shorter-shifted *SWS* genes in amphibious species more generally.

Overall, opsin orthologs were highly conserved between Salariini species, and only very few changes in key tuning sites potentially shifting spectral sensitivities could be found (Table 1 and Table S3). Therefore, neither at the opsin-sequence nor at the expression level were there any apparent adaptations to the transition from water to land. This is surprising since morphological adaptations of the visual system when moving out-of-water are common (Sayer 2005). It remains possible that instead of relying on their opsins, Salariini blennies change the chromophore part of the photopigment. For example, most amphibians use red-shifted chromophore A2 based visual systems during their aquatic life stages, but change to a blue-shifted A1 based visual system after metamorphosis and moving onto land (Wilt 1959, Liebman and Entine 1968, Bridges 1972). Most Salariini species were found to lowly express *Cyp27c1*, the enzyme responsible for the A_1_ to A_2_ switch (Enright et al. 2015) (Table 2). It remains to be seen if the use of these chromophores differs between habitats.

In summary, we found no apparent molecular adaptations in terms of opsin gene sequence variability and expression in terrestrial blennies when compared to their amphibious and aquatic sister species. On the contrary, our data suggests that Salariini blennies evolved visual systems early on that were ideal to conquer shallow reef and intertidal habitats, and no further molecular adaptations have been made for life on land.

## Acknowledgements

We would like to thank Sara Stieb and Fanny de Busserolles for assistance with RNA extractions and specimen collection, and Janette Edson from the Queensland Brain Institute’s sequencing facility for library preparation and RNA sequencing.

## Author Contributions

T.J.O. conceived the study and designed the experiments together with F.C. and K.L.C. T.J.O., G.M.C. and F.C. collected the specimens. F.C. performed the experiments, analysed the data, and wrote the initial manuscript. All authors reviewed and approved the final version of the manuscript.

## Data Accessibility

Raw-read transcriptomes (SRA tba) and single gene sequences (#tba) are available through GenBank (https://www.ncbi.nlm.nih.gov/genbank/). Gene alignments and single gene phylogenies can be accessed through Dryad (#tba). All other data is given either in the main manuscript or the supplementary material.

